# The Sez6 family inhibits complement at the level of the C3 convertase

**DOI:** 10.1101/2020.09.11.292623

**Authors:** Wen Q. Qiu, Shaopeiwen Luo, Stefanie A. Ma, Priyanka Saminathan, Herman Li, Jenny Gunnersen, Harris A. Gelbard, Jennetta W. Hammond

## Abstract

The Sez6 family consists of Sez6, Sez6L, and Sez6L2. Its members are expressed throughout the brain and have been shown to influence synapse numbers and dendritic morphology. They are also linked to various neurological and psychiatric disorders. All Sez6 family members contain 2-3 CUB domains and 5 complement control protein (CCP) domains, suggesting that they may be involved in complement regulation. We show that all Sez6 family members inhibit C3 deposition by the classical and alterative pathways with varying degrees of efficacy. For the classical pathway, Sez6 is a strong inhibitor, Sez6L2 is a moderate inhibitor, and Sez6L is a weak inhibitor. Using Sez6L2 as the representative family member, we show that it specifically deactivates C3 convertases by accelerating the decay or dissociation of the C3 convertase components. Sez6L2 also deactivates C3 convertases of the alternative pathway by serving as a cofactor for Factor I to facilitate the cleavage of C3b. However, Sez6L2 has no cofactor activity toward C4b. In summary, the Sez6 family are novel complement regulators that inhibit C3 convertases.

## Introduction

The Sez6 family consisting of Sez6, Sez6L, and Sez6L2, is notable because its members have been identified as potential susceptibility genes for multiple neurodevelopmental and psychiatric disorders including: autism, schizophrenia, intellectual disability, epilepsy, and bipolar disorder [1–9]. Sez6L2 is located in the 16p11.2 deletion/duplication region which encompasses 26 genes and accounts for ~1% of all ASD cases but also has strong links to schizophrenia and intellectual disability [3, 4, 10]. Sez6L2 is also part of a smaller 16p11.2 deletion region identified in a family with ASD that spans only 5 genes [11]. Sez6 family proteins are expressed by neurons throughout the brain during development and in adulthood. Some brain regions express multiple Sez6 family members, suggesting possible redundancy, while other regions have differential expression [12–16]. In experimental mouse models, the Sez6 family has been shown to modulate synapse numbers, synaptic plasticity, and dendrite morphology in the cortex and hippocampus and neuronal connectivity in the cerebellum. Genetic loss of Sez6 genes results in impaired cognition, motor learning, and motor functions [14–17]. Outside the nervous system, Sez6 family members have been implicated as markers of poor prognosis in cancer [18–24]. Although a few binding partners of the Sez6 family have been proposed [25–29], the molecular mechanisms and functions of Sez6 proteins are still unclear.

The domain structure of Sez6 family members consisting of 5 complement control protein (CCP) domains and 2-3 CUB domains, suggest they may have activity connected to the complement cascade. CCP domains, also known as short consensus repeats or SUSHI repeats, contain ~ 60-70 amino acids with four invariant cysteines forming disulfide bonds. CCP domains are the primary components of several proteins of the complement system [30]. However, CCP domains are also found in a variety of proteins outside the complement pathway involved in cell adhesion, blood coagulation, and signal transduction related to cytokines and neurotransmission. CUB domains are named after their founding members: complement <u>C</u>1r/C1s, <u>u</u>EGF, and <u>B</u>MP1 and primarily mediate protein-protein interactions. CUB domains are found in complement proteins such as C1s, C1r, MASP-1/2, and C3 as well as many other non-complement pathway proteins. This goal of this study was to evaluate the Sez6 family for complement regulatory functions.

The complement system has three initiating pathways (classical, alternative, and lectin) that all create C3 convertases to cleave C3 into C3a and C3b. C3b is an opsonin that facilitates phagocytosis and is deposited onto activating surfaces such as antibody complexes, cellular debris, foreign particles, or even an organism’s own healthy “self” tissue. C3b is also a central component of C3 and C5 convertases and when significantly accumulated it can initiate the lytic complement pathway leading to cell lysis. Complement activation on self cells is largely controlled by the regulators of complement activation (RCA) protein family that protect “self” surfaces by limiting the activity of the C3 convertases (C3bBb or C4b2a). RCAs bind C3b or C4b, then inactivate the C3 convertases by either removing the active protease or by recruiting the cleaving enzyme, complement Factor I, to destroy C3b or C4b [30–32]. Cleaved C3b fragments (iC3b, C3c, C3dg/d) can no longer participate in the complement cascade leading to cell lysis, but they still have opsonizing and immune signaling properties.

The complement system is a fundamental part of the innate immune system, but it also has important roles in neural development including neurogenesis, synapse pruning, and neuronal migration [33–38]. It additionally mediates immune responses to prenatal or early postnatal brain insults that affect brain development with lifelong consequences (reviewed in [38]). Furthermore, complement contributes to pathological cell and synapse loss in aging and diseases including Alzheimer’s, viral encephalitis, glaucoma, lupus, epilepsy, schizophrenia, frontotemporal dementia, and multiple sclerosis [33–35, 39–52]. Therefore, understanding complement regulation in the brain is important for neurodevelopment as well as neurodegenerative disease. We show here that members of the Sez6 family, are novel complement regulators that inhibit the complement pathway at the level of the C3 convertases.

## Results

### Expression of Sez6L2 and other RCAs in hippocampal neurons

Self-directed complement activity is usually tightly controlled by complement regulatory proteins expressed on cell membranes and/or secreted in soluble form. However, the identity of complement regulators functioning on neurons throughout neuronal development or in disease with a potential role in modulating complement-mediated synaptic pruning has yet to be fully elucidated. A common feature of complement regulators, particularly those that work at the level of C3 convertases, is that they contain three or more tandem CCP domains [30, 32]. Because hippocampal neurons express only trace amounts of the most well-known RCAs (DAF, CR1/Crry, Factor H (FH), C4BP, and MCP) or C1-INH, a complement regulator that is not part of the RCA family [53] (Figure 1A), we searched the smart protein database (smart.embl-heidelberg.de/) for proteins with at least three tandem CCP domains that are also highly expressed by neurons in development. We identified the Sez6 family of proteins (consisting of Sez6, Sez6L, and Sez6L2) as ideal candidates. RNA-Seq expression data from the Hipposeq database (http://hipposeq.janelia.org; [53]) show that the Sez6 family members are highly expressed by pyramidal neurons in the hippocampus with differential expression in CA1, CA3, and dentate gyrus (Figure 1A). Immunostaining of adult mouse brain sections shows the Sez6L2 protein is localized to neuronal cell bodies of the CA1 region, but also has significant co-localization with synapses (identified as Homer1+ puncta) in the stratum radiatum (Figure 1B-C).

**Figure 1:**
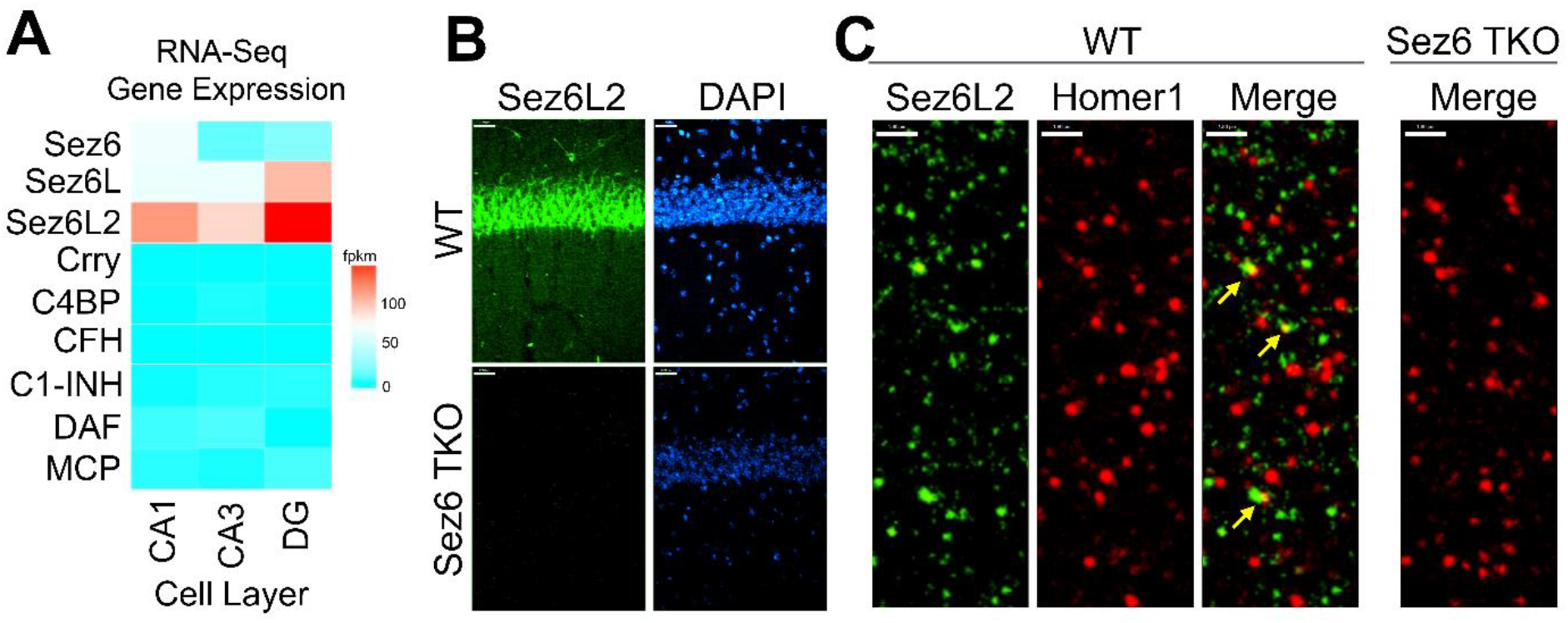
Sez6 family expression in the hippocampus. A) Sez6, Sez6L, and Sez6L2 are expressed by principal (excitatory, pyramidal) neurons of the mouse hippocampus at much higher levels than other known complement regulators (namely Crry, C4BP, CFH, C1-INH, DAF, and MCP). Expression data was obtained from Hipposeq: a comprehensive RNA-Seq database of gene expression in hippocampal principal neurons (http://hipposeq.janelia.org; [53]). The RNA samples used in this database were isolated from mouse hippocampal principal neurons micro-dissected from the CA1, CA3, or Dentate Gyrus (DG) cell layers of the hippocampus at Postnatal Day 25-32. Differential gene expression is shown in the heatmap with the relative units of FPKM (Fragments Per Kilobase of Exon Per Million Reads Mapped.) B) Brain sections from adult WT mice or Sez6 triple knockout mice (TKO) were immuno-stained for Sez6L2 (green) and DAPI and imaged in the CA1 region of the hippocampus. Scale Bar= 27μm. High density Sez6L2 staining occurs around cell bodies in the pyramidal layer, but significant Sez6L2 is also found in the stratum radiatum and stratum oriens. C) Higher magnification images of sections immuno-stained for Sez6L2 (green) and the postsynaptic protein, Homer1 (red), shows Sez6L2 is found near or co-localized with synapses in the stratum radiatum. Scale bar = 1.8μm.

### Recombinant Sez6L2-MH is a soluble, truncated form of Sez6L2 with a Myc-6xHis tag

After having identified the Sez6 family proteins as potential complement regulators due to their domain structure, we sought experimental evidence to support this putative function. As many traditional complement assays use soluble proteins, we first generated a truncated expression construct of human Sez6L2, named Sez6L2-MH, which replaced the transmembrane domain and cytoplasmic tail with a tandem Myc-6xHis tag. This Sez6L2-MH plasmid was expressed in HEK293 cells grown in serum free media and the secreted protein was purified via the His-tag. The purity and identity of the protein was verified by SDS-PAGE under reducing conditions followed by Coomassie staining and anti-Sez6L2 or anti-Myc western blots (Figure 2A-B).

**Figure 2:**
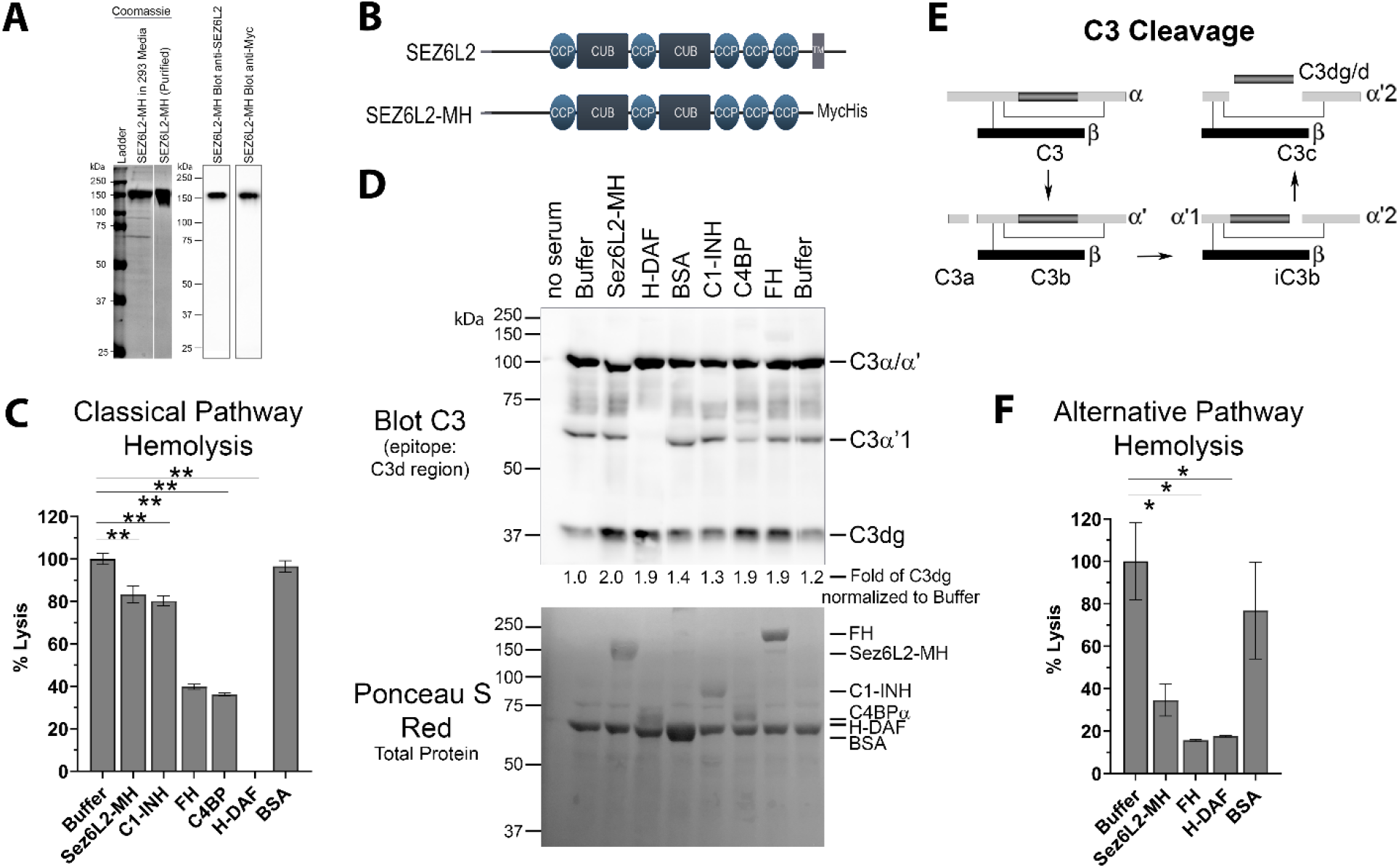
Sez6L2 prevents classical and alternative pathway hemolysis. A) Purified Sez6L2-MH is shown by a coomassie stained gel and by western blot with anti-Sez6L2 and anti-Myc antibodies. Lanes with the coomassie stain are from the same gel. B) Schematic of Sez6L2 and Sez6L2-MH domain structures. CCP=Domain abundant in <u>c</u>omplement <u>c</u>ontrol <u>p</u>roteins. CCP domains are also known as SUSHI repeats or short complement-like repeat (SCR) CUB= Domains named after complement <u>C</u>1r/C1s, <u>u</u>EGF, and <u>B</u>MP1 TM=Transmembrane region. Sez6L2-MH was made by replacing the transmembrane and cytoplasmic tail domains with a tandem Myc, 6xHis tag. C) and D) Classical pathway hemolysis assay. Antibody-coated sheep erythrocytes were exposed to human serum pre-incubated with purified Sez6L2-MH, C1-INH, FH, C4BP, H-DAF, or BSA. After 30 mins, the percent of cell lysis was measured by spectrophotometry (A415). C1-INH, FH, C4BP, and H-DAF are known complement inhibitors and were used as positive controls. BSA was used as a negative control for comparison. H-DAF is His-tagged DAF. 1-way ANOVA (P <0.0001; F(6,14)=314.4). N=3 (1 experiment with 3 replicates; Representative of 4 independent experiments with Sez6L2 and buffer and 2-3 experiments with each control protein). D) Supernatants from classical pathway hemolysis assays were further analyzed for C3 cleavage products by western blot. Supernatants in D are from same experiment shown in C. Sez6L2-MH increases the amount of the C3dg cleavage product similar to other complement regulators that work at the level of the C3 convertase. Sez6L2-MH runs just above the C3α band of C3. When large amounts of Sez6L2 are present it can cause C3α to run lower on the gel and sometimes mildly interferes with antibody binding to C3α. The membrane blotted for C3 was also stained with ponceau S to reveal total protein and shows the presence of the purified proteins in each sample. E) Schematic of C3 cleavage products. The C3d region recognized by our C3 antibody is highlighted in dark grey. F) Alternative pathway hemolysis assay. Rabbit erythrocytes were exposed to human serum pre-incubated with Sez6L2-MH, complement regulators, or BSA in presence of 12.5mM MgEGTA to block the classical pathway. Then the percent of cell lysis was measured by spectrophotometry (A415). 1-way ANOVA (P =0.0122; F(4,7)=7.298). Results are the mean of three independent experiments which tested 4 independent Sez6L2-MH preps multiple times. N=3 for buffer and Sez6L2-MH; N=2 for FH, H-DAF, and BSA. For all graphs * = p <0.05; ** = p<0.01

### Sez6L2 Inhibits Classical and Alternative Pathway Hemolysis

Complement activation by the classical and alternative pathways ultimately results in the formation of the membrane attack complex which directly lyses target cells. As such, complement-mediated lysis of erythrocytes, also known as hemolysis, is a useful assay to measure the ability of Sez6L2-MH to regulate the complement pathway. We tested Sez6L2-MH alongside complement inhibitory proteins C1-INH, FH, and C4BP purified from human serum and His-tagged DAF (H-DAF) purified similarly to Sez6L2-MH. BSA was also tested as a negative control protein. Sez6L2-MH was able to modestly inhibit complement-mediated lysis of sheep erythrocytes by the classical pathway, exhibiting 83 ± 4% lysis compared to 100 ± 3% lysis by buffer alone (p<0.001; Holm-Sidak MCT; Figure 2C). Sez6L2-MH performed at a similar level as C1-INH (80 ± 2% lysis), but not as well as FH, C4BP, or H-DAF (40 ± 1%, 36 ± 1% or 0 ± 0% lysis respectively). As expected, BSA had no effect on erythrocyte lysis by the classical pathway. Interestingly, when we assayed the supernatants from classical pathway hemolysis experiments, we found that Sez6L2-MH samples generated the C3 cleavage product, C3dg, at a similar level as all the complement inhibitors that operate at the level of C3 convertases, specifically, H-DAF, FH, and C4BP (Figure 2D-E). This suggests that Sez6L2’s mechanism of action may be more related to the functions of these regulators than the mechanisms of C1-INH.

A similar hemolysis assay was used for the alternative pathway, but was performed in the presence of EGTA. C1 of the classical pathway requires calcium which is preferentially chelated by EGTA, effectively preventing classical pathway initiation while sparing the alternative pathway. Sez6L2-MH was able to significantly inhibit RBC lysis by the alternative pathway, showing 34 ± 8% lysis compared to 100 ± 18% lysis by buffer alone (p<0.001). Comparatively, FH limited lysis to 16 ± 0.5% and H-DAF to 18 ± 0.5% (Figure 2F). Thus, Sez6L2 is an inhibitor of the classical and alternative complement pathways *in vitro*.

### Sez6L2 is a cofactor for Factor I (FI) cleavage of C3b but not C4b

As the classical pathway hemolysis supernatants suggested that Sez6L2 enhanced the formation of the C3dg, we next tested whether Sez6L2 functions as a cofactor for FI to facilitate cleavage of C3b or C4b. FH and C4BP are known co-factors of FI towards C3b and/or C4b and were used as positive controls. C3b is composed of the C3α’ chain and the C3β chain linked by disulfide bonds (Figure 3A). Incubation of C3b and FI with 1 μg FH or C4BP for 2 hours resulted in an almost complete loss of C3α’ and the appearance of the two cleavage products α’1 (visible by western blot with a C3d antibody and in the Coomassie stained gel) and α’2 (visible in the Coomassie stained gel only) (Figure 3A-B). FI cleavage of C3b with 1 μg of Sez6L2-MH resulted in only a small amount of C3b cleavage, but higher amounts of Sez6L2 (2μg-10μg) facilitated FI cleavage of the C3α’ chain to α’1 and α’2 in a dose-dependent manner. We also found that increased incubation times of 4 or 8 hours increased the amount of C3α’ cut by FI in the presence of Sez6L2-MH (Supplemental figure 2A). Sez6L2 did not direct FI to make a second cut in the C3α’ chain to generate C3dg in this assay. This second FI cleavage site seems to be primarily utilized by the CR1 cofactor and perhaps other proteases [31, 54]. FH, but not C4BP, facilitated a small amount of cleavage at the additional site to generate C3dg. Incubation with Sez6L2 and C3b alone did not result in any C3b cleavage products, indicating that FI is the active protease responsible for C3b cleavage in the presence of Sez6L2-MH. Additionally FI by itself is unable to cut C3b.

**Figure 3:**
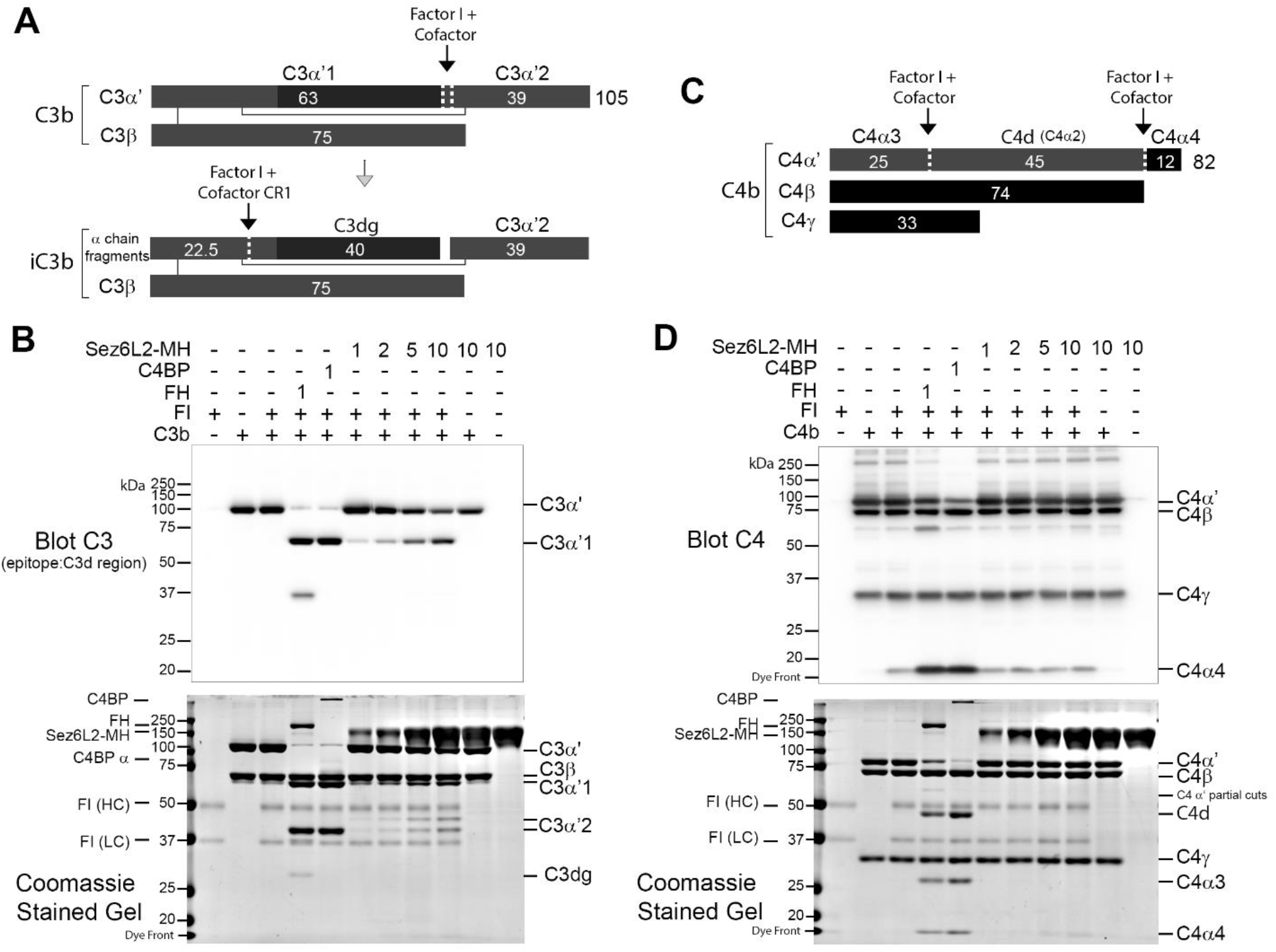
Sez6L2 acts as a cofactor for Factor I cleavage of C3b but not C4b. A) Schematic of Factor I and cofactor cleavage of C3b and iC3b. B) Factor I cleavage assay of C3b. C3b and Factor I (FI) were incubated alone, with Factor H (FH), or C4BP, or with concentrations of Sez6L2-MH ranging from 1 to 10 μg/mL for 2 hours at 37°C. Then samples were analyzed by western blot using antibodies that recognize C3d, a region within the C3α chain (and highlighted by the black rectangle in the schematics in (A)). Coomassie stained gels are also shown. FH and C4BP are known co-factors of FI towards C3b and served as positive controls. Incubation of C3b and FI with Sez6L2-MH also generated the C3 cleavage products α’1 and α’2 showing Sez6L2-MH is a cofactor for Factor I cleavage of C3b. C) Schematic of Factor I + cofactor cleavage of C4b. D) C4b and Factor I (FI) were incubated alone, with FH, C4BP, or with concentrations of Sez6L2-MH ranging from 1 to 10 μg/mL for 2 hours at 37°C. Then samples were analyzed by western blot using a C4 polyclonal antibody or coomassie stained gels. C4 components recognized by the C4 antibody are colored black in the schematic in C. C4BP is a known cofactor of FI for C4b cleavage and served as a positive control. Incubation of C4b and FI with Sez6L2-MH did not result in the appearance of C4b cleavage products.

Some FI cofactors are specific for either C3b or C4b, while others work on both. Therefore, the FI cleavage assay was repeated with C4b. C4b is composed of 3 peptide chains C4α’, C4β, and C4γ attached by disulfide bonds. Cofactors aid FI to cut the C4α’ chain at two locations yielding C4d, C4α3, C4α4 (Figure 3C). C4b and FI were incubated alone, with 1μg of FH or C4BP, or with concentrations of Sez6L2-MH ranging from 1 to 10 μg/mL for 1-2 hours. C4BP was able to facilitate the almost complete cleavage of C4α’ to C4α3, C4α4, and C4d while FH facilitated some partial and low level cleavage. No amount of Sez6L2-MH tested was able to promote any FI cleavage of C4b (Figure 3D). Longer incubation times of 4 or 8 hours also did not allow Sez6L2-MH to promote FI cleavage of C4b (Supplemental Figure 2B) Therefore, Sez6L2 is a cofactor for FI, facilitating cleavage of C3b, but not C4b. This means Sez6L2 can help breakdown the alternative pathway C3 convertase (C3bBb) but not the classical and lectin pathway C3 convertases (C4b2a). Yet, cleavage of C3b, can still reduce the downstream impact of the classical pathway, by limiting the formation or lifespan of the C5 convertase (C4b2aC3b) and restricting the amplification of the classical pathway provided by the alternative pathway.

### Sez6L2 accelerates the decay of C3 convertases

Complement inhibitors that act at the level of the C3 convertase not only function as cofactors for FI, but they can also act as decay accelerating factors to permanently remove the activated protease component of the C3 convertase from the complex (Bb is removed from the alternative convertase C3bBb; 2a is removed from the classical/lectin convertase, C4b2a). MCP functions primarily as a cofactor and does not have decay accelerating activity. Alternatively, DAF was named for its decay accelerating activity and has no cofactor function. FH has both cofactor and decay accelerating activity [31]. To find out if Sez6L2-MH has decay accelerating activity for the alternative pathway C3 convertase, we coated C3b on a 96 well plate and then added Factor B and Factor D to generate C3bBb bound to the plate. FH and Sez6L2-MH were added to the wells in increasing concentrations. The ability of Sez6L2-MH or FH to displace Bb from the bound C3bBb complex was assessed in an ELISA assay using an anti-FB antibody. The percent convertase remaining on the plate (normalized to a buffer only control) was graphed against the concentrations of Factor H and Sez6L2-MH. Sez6L2-MH at lower concentrations (0.01 μg/mL - 5 μg/mL) had no effect on the amount of intact C3 convertase, while higher concentrations of Sez6L2 (50 μg/mL – 500 μg/mL) significantly decreased the amount of intact C3 convertase by 40-80% in a dose dependent manner (Figure 4A). FH was effective at much lower concentrations, as 0.05 μg/mL - 5 μg/mL significantly decreased the amount of C3 convertase by 20-75%. Higher concentrations of FH (25 μg/mL – 200 μg/mL) were also effective and showed a plateauing effect and reduced the convertase by just over 75%. The relative IC50 for FH was 0.0102 μg/mL and the relative IC50 for Sez6L2-MH was 87.41 μg/mL. Supernatants collected from the plate at the end of the incubation with Sez6L2-MH or FH were collected and analyzed by western blot with antibodies to Factor B. With increasing levels of Sez6L2-MH or FH, we found increasing levels of Bb (MW 70 KDa) in the supernatant, which mirror the deceasing amounts of Factor B bound to the ELISA plates (Figure 4B). Taken together, these data show that Sez6L2-MH exhibits decay accelerating activity for the alternative pathway C3 convertase C3bBb, but requires higher concentrations than FH.

**Figure 4:**
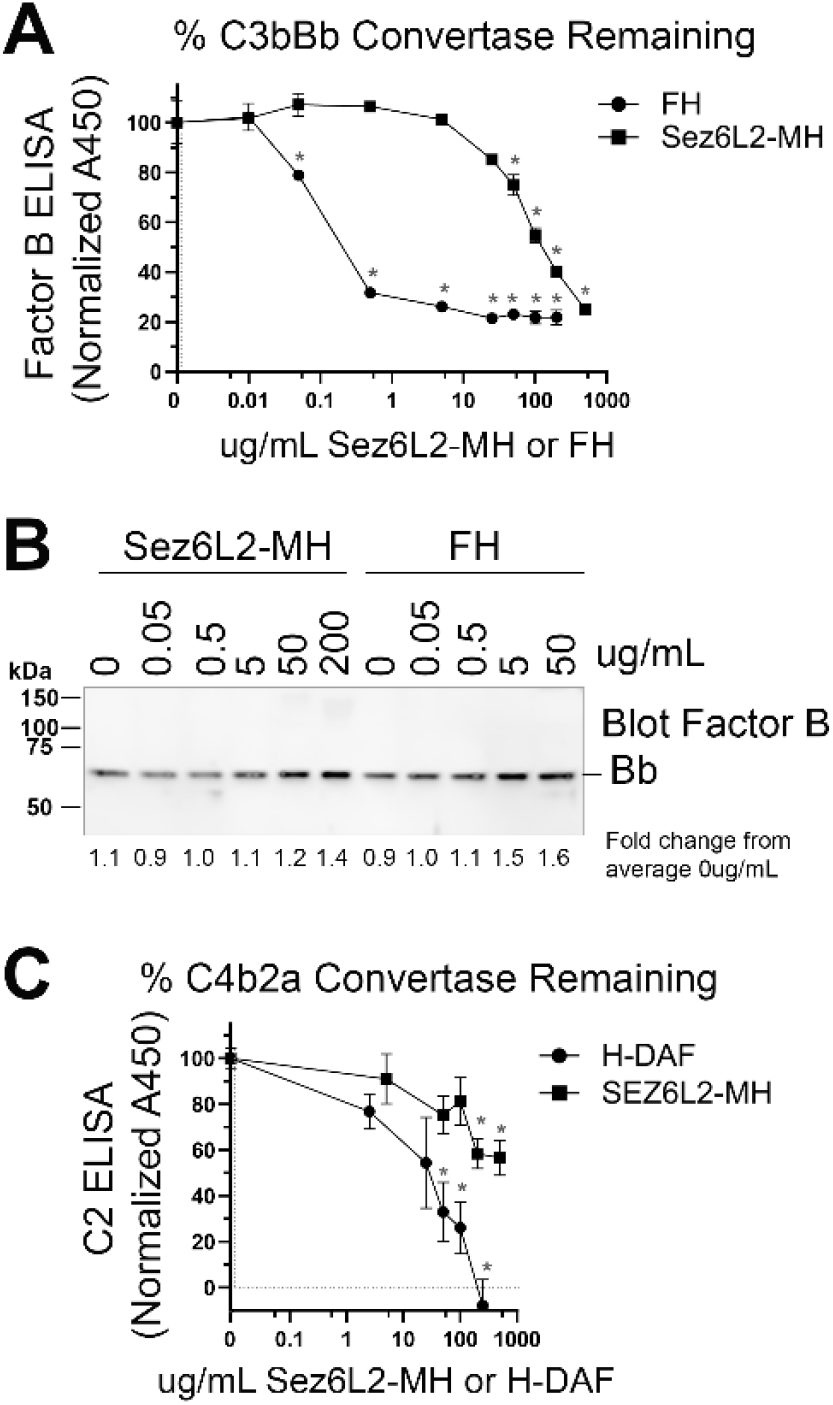
Sez6L2 has decay accelerating activity for the alternative and classical/lectin pathway C3 convertases. A and B) Alternative C3 convertase assay: A 96 well plate coated with C3b was incubated with Factor B and Factor D to form the C3 convertase C3bBb, then incubated with Sez6L2-MH or FH at concentrations ranging from 0 to 500 μg/mL to assess their decay accelerating activity. Factor B remaining bound to the plate (as C3bBb) was detected using an anti-Factor B antibody ELISA in (A) and Bb released into the supernatant is shown via western blot (B). C) Classical C3 convertase assay: A plate coated with C4b was incubated with C2 and C1s-enzyme to form the classical/lectin pathway C3 convertase C4b2a, then incubated with Sez6L2-MH or H-DAF at concentrations ranging from 0 to 500 μg/mL to assess their decay accelerating activity. C2 remaining bound to the plate (presumably as C4b2a) was detected using an anti-C2 antibody ELISA. For A and C ELISAs: N=3 (1 experiment with 3 replicates; representative of 2-3 independent experiments). Statistics: 1-way ANOVAS with Holms-Sidak multiple comparison’s tests were performed for each complement inhibitor with comparisons to 0 ug/mL controls. * p<0.05.

Next we tested whether Sez6L2-MH could accelerate the decay of the classical/lectin pathway C3 convertase, C4b2a. A plate coated with C4b was incubated with C2 and C1s-enzyme to form the C3 convertase, C4b2a, then incubated with Sez6L2-MH or H-DAF at concentrations ranging from 0 to 500 μg/mL to assess their decay accelerating activity. C2 remaining bound to the plate (presumably as C4b2a) was detected using an anti-C2 antibody ELISA (Figure 4C). H-DAF effectively removed bound C2a from the plate in a dose dependent manner with 250ug/mL H-DAF resulting in complete dissociation of C2a. Sez6L2-MH was also able to dissociate the C2a in a dose dependent manner with the maximal dose of 500ug/mL Sez6L2-MH reducing C2a by 43% (plate bound C2a = 57 ± 8% % vs 100 ± 4% % with buffer alone; p=0.002 Holm-Sidak MCT). Thus Sez6L2-MH has decay accelerating activity for the classical/lectin pathway C3 convertase as well as the alternative C3 convertase, but is less effective than H-DAF.

### Full length Sez6L2, Sez6, and Sez6L inhibit complement deposition on CHO cells

The truncated Sez6L2-MH purified protein was very useful in examining the specific mechanisms by which Sez6L2 inhibits complement at the level of the C3 convertase. However, we also found it was less effective than the positive control, soluble complement regulators from human serum. It is possible that the purification process leads to partial deactivation of Sez6L2-MH, necessitating higher levels of Sez6L2-MH to achieve complement inhibition. It is also possible, that the truncated version could be less active than the full-length transmembrane protein due to its inability to localize to areas of complement activation. Naturally soluble, serum regulators like FH, C4BP, and CR1 have multiple C3 binding sites or can oligomerize to enhance homing to areas of active complement deposition. They also have binding sites for other complement factors or “self” membranes that aid complement inhibition and protection of “self” surfaces. These properties may also help explain their superior activity compared to Sez6L2-MH. Yet, this also means the activity of truncated complement inhibitors, like FH, CR1 or SUSD4 does not always equal the activity of the entire protein [31, 55, 56]. Thus, it is important to test the inhibitory action of full length Sez6L2 on cell surfaces.

First, CHO (Chinese hamster ovary) cells were transfected with GFP alone or co-transfected with GFP and Myc-tagged human Sez6L2 (M-Sez6L2) or His-tagged DAF (H-DAF). The cells were coated in antibodies from anti-hamster serum to help initiate the classical pathway. Then the cells were exposed to C5-depleted human serum ranging in concentration from 0-20%. C5-depleted serum was used to prevent cell lysis, but still allow full C3 deposition which was detected with antibodies to C3b/C3c. GFP positive cells co-transfected with M-Sez6L2 showed ~50% reduced complement deposition at all serum levels compared to cells transfected with GFP alone (at 15% serum; p=0.007, Holm-Sidak MCT) (Figure 5A-B). Cells expressing H-DAF, reduced C3 deposition to ~13-15% compared to GFP alone (at 15% serum p=0.001).

**Figure 5:**
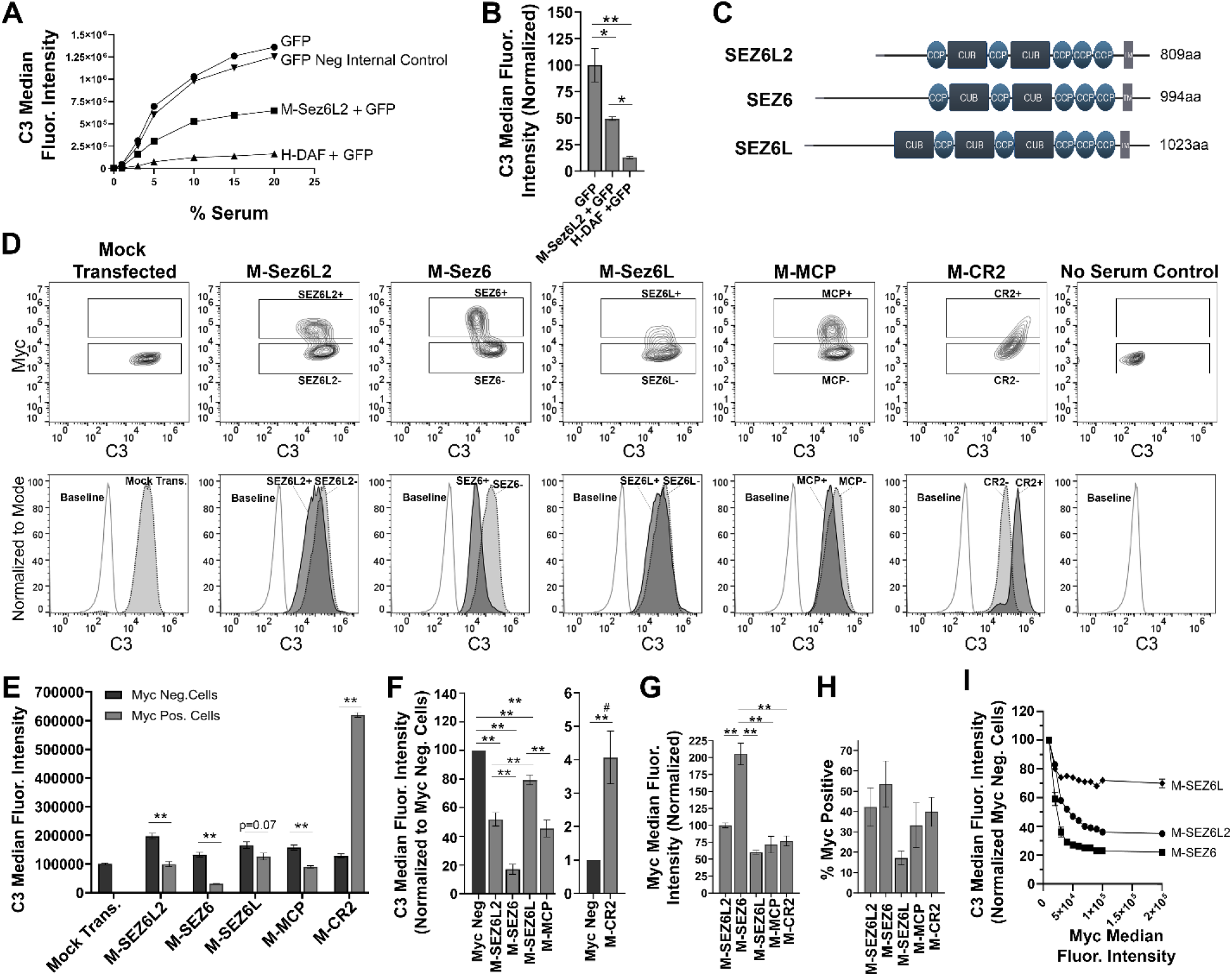
Full Length Sez6L2, Sez6, and Sez6L inhibits C3 deposition on CHO cells by the classical pathway. A-B) Sez6L2 inhibits C3 deposition at a range of serum concentrations. CHO cells were transfected with plasmids for GFP alone or with Myc-tagged Sez6L2 (M-Sez6L2) or His-tagged DAF (H-DAF). CHO cells were coated with antibodies and exposed to 0-20% C5 depleted human serum for 1 hour and then labeled with anti-C3b/C3c antibodies and analyzed by flow cytometry. One experiment is shown that is representative of two independent experiments. B) C3 deposition on GFP transfected cells with or without M-Sez6L2 or H-DAF at 15% serum. ANOVA (P=0.0016; F(2,6)=22.51). N=3; 1 experiment with 3 replicates (representative of 3+ independent experiments). C) Schematic of Sez6L2, Sez6, and Sez6L protein domain structures. D-I) CHO cells were transfected with the indicated Myc-tagged cDNAs and processed as outlined in A with 15% C5 depleted serum, except that an anti-Myc antibody was used in place of GFP to identify transfected and expressing CHO cells. D) 5% Contour plots of C3 versus Myc fluorescence (top layer) and C3 fluorescence histograms (bottom layer) of the same samples normalized to mode and compared to baseline cells not exposed to serum. For Contour plots, boxed regions highlight cells designated as myc positive (top box) and myc negative (lower box) populations. For C3 histograms, dark grey, solid line population = Myc positive cells; Light grey, dotted line population= Myc negative cells; White, dashed grey line population = baseline. Representative of 4+ independent experiments. E) Quantification of the average median C3 fluorescence intensity from myc positive and myc negative cells within each sample. Statistics = t-tests. N=3 (1 experiment with 3 replicates; Representative of 4+ independent experiments). F) Average median C3 fluorescence intensities after normalization to the myc negative cells from each experimental group. ANOVA between Myc+ cell populations (p<0.001; F(4, 15)=64.53). Sez6L2 inhibits C3 deposition at a level comparable to positive control MCP. Sez6 is a stronger complement inhibitor than Sez6L2 and Sez6L is a weaker inhibitor. F) Average median Myc fluorescence intensity from myc positive cells ANOVA (p<0.001; F(4, 15)=36.79). G) Average % of Myc positive cells in each experimental group (ANOVA, p=0.115; F(4, 15)=2.224). For sections F-H, N=4 (4 independent experiments). I) Sez6 blocks complement deposition more efficiently than Sez6L2 and Sez6L even when comparing similar levels of myc surface expression. Average C3 median fluorescence intensity normalized to internal myc negative populations for M-Sez6, M-Sez6L2, and M-Sez6L samples shown relative to the myc median fluorescence intensity. N=3 (1 experiment with 3 replicates, Representative of 3 independent experiments). For all graphs * = p <0.05; ** = p<0.01; # = p<0.001 for all Myc+ groups compared to M-CR2.

Next we compared the complement inhibitory activity of full length Myc-tagged Sez6L2 to its family members Sez6 and Sez6L (which are 49% and 50% identical to Sez6L2 respectively; Figure 5C) to learn whether complement inhibition is a family function or limited to Sez6L2. We also compared the Sez6 family members to two CCP domain containing proteins, Myc-tagged MCP or CR2. MCP is a relatively small transmembrane complement inhibitor composed of just 4 tandem CCP domains. CR2 is also a transmembrane protein composed of 15-16 tandem CCP domains and expressed by immune cells. CR2 is not a complement inhibitor, but instead binds C3d/C3dg to present complement opsonized antigens to the adaptive immune system as well as initiate signal transduction cascades [57]. It can also enhance C3b deposition by the alternative pathway on expressing cells [58, 59] and serves as a control in this paradigm for what excessive complement deposition looks like. Mock transfected cells and cells without serum exposure were used as controls for full complement activation or a negative baseline respectively. First we assayed complement deposition initiated primarily by the classical pathway. M-Sez6L2, M-Sez6, M-Sez6L, and M-MCP all significantly inhibited C3 complement deposition and M-CR2 significantly increased deposition compared to non-transfected, Myc negative cells within each experimental sample (Figure 5D-F). Overall, M-Sez6 was the most effective at decreasing complement deposition with Myc positive cells having only 17 ± 3% of the level C3 found on internal Myc negative cells (p<0.001, Holm-Sidak MCT). M-Sez6L2 expressing cells had 52 ± 5% the deposition of internal Myc negative cells (p<0.001) M-Sez6L had 79 ± 3% (p=0.004) and M-MCP had 45 ± 6% (p<0.001). M-CR2 on the other hand, had 407 ± 78% the deposition of internal Myc negative cells p<0.001) (Figure 5F). These results show that Sez6 family members share the complement inhibitory function but have different levels of activity. Sez6 is the most effective inhibitor of the classical pathway and functions at a level equivalent to H-DAF. Sez6L2 is a moderate inhibitor and functions at a level comparable to MCP. Sez6L is a weak inhibitor.

Because M-Sez6 expressed on the cell surface at twice the level of Sez6L2, Sez6L, and MCP (Figure 5G) but usually in a similar percentage of cells (Figure 5H), we wondered whether M-Sez6 was a more effective inhibitor simply because of the higher expression levels or whether there are intrinsic activity differences in the family members. Thus we gated the cell populations based on increasing levels of Myc surface expression to compare complement deposition levels in cells with similar levels of surface M-Sez6, M-Sez6L or M-Sez6L2. The results suggest that all Sez6 family members inhibit complement better with higher surface expression. However, M-Sez6 was still more effective at limiting C3 complement deposition than M-Sez6L and M-Sez6L2 at all expression levels (Figure 5I).

Finally, full-length Sez6L2, Sez6, and Sez6L were analyzed for their ability to inhibit complement deposition initiated primarily by the alternative pathway in the presence of Mg-EGTA. The results were similar to that obtained with the classical pathway and showed all Sez6 family members to be effective inhibitors of C3 deposition. However, their activities relative to each other were different (Figure 6A-C). M-Sez6L2 expressing cells had 34 ± 1%, M-Sez6 had 59 ± 2%, M-Sez6L had 52 ± 6%, and M-MCP had 65 ± 2% and M-CR2 had 2624 ± 256% the deposition of internal Myc negative cells (Figure 6C). C3 deposition by the alternative pathway was less intense than the classical pathway and resulted in C3b/C3c coated cells segregating into two main population peaks. The primary peak population had low C3 deposition. The second, smaller peak population, had high C3 deposition at similar levels as found in the classical pathway assays. M-Sez6L2, M-Sez6, Sez6L, and M-MCP all inhibited C3 complement deposition by limiting expressing cells from reaching the complement deposition levels of the high C3b peak population (Figure 6A). Alternatively, M-CR2 increased C3 deposition, shifting all Myc positive cells into the high C3 population. Interestingly, M-Sez6L2 expressing cells also had decreased C3 levels in the lower peak population relative to the non-transfected, Myc negative cells within the same sample (Figure 6A,B). However, these Myc negative cells in M-Sez6L2 samples often had increased C3 deposition compared with the Myc negative cells in other samples raising the question of whether M-Sez6L2 protects expressing cells at the expense of promoting complement deposition on non-expressing cells. Nevertheless, the cells expressing Sez6 family members had less complement deposition by the alternative pathway than non-expressing cells and their complement inhibitory activity equaled or exceeded the activity of the known complement regulator, MCP. In summary, Sez6 family members are effective inhibitors of C3 complement deposition by both the classical and alternative pathways but individual Sez6 family members vary in the efficacy of their complement inhibitory activity toward each pathway.

**Figure 6:**
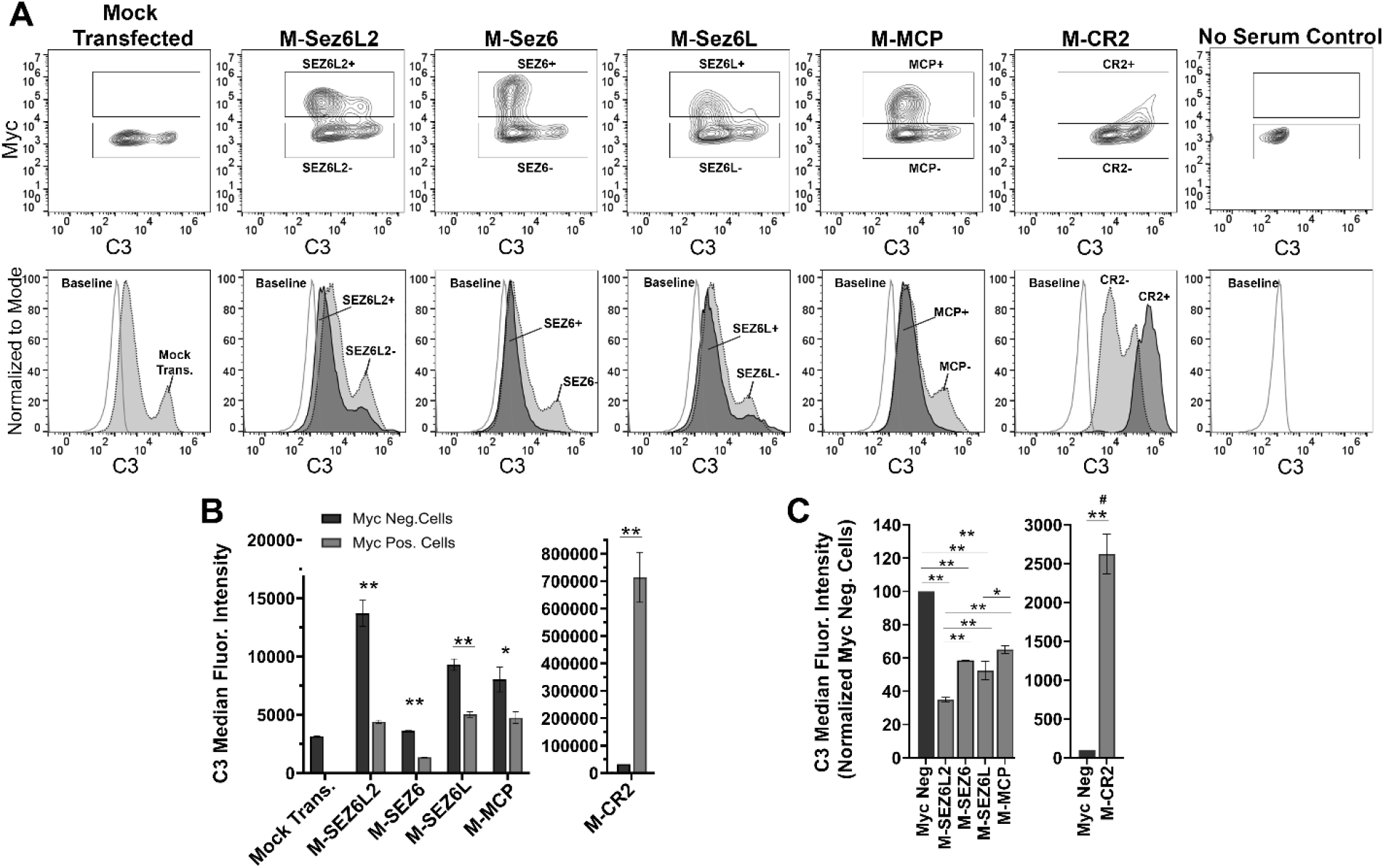
Full Length Sez6L2, Sez6, and Sez6L inhibit C3 deposition on CHO cells by the alternative pathway. CHO cells were transfected with the indicated Myc-tagged cDNAs and then coated with a low level of antibodies and exposed to 15% C5 depleted human serum for 1 hour in the presence of 10mM EGTA and 10mM MgCl_2_ to block the classical pathway. Cells were then labeled with anti-C3b/C3c and anti-Myc antibodies and analyzed by flow cytometry. A) 5% Contour plots of C3 versus Myc fluorescence (top layer) and C3 fluorescence histograms (bottom layer) of the same samples normalized to mode and compared to baseline cells not exposed to serum. For Contour plots, boxed regions highlight cells designated as myc positive (top box) and myc negative (lower box) populations. For C3 histograms, dark grey, solid line population = Myc positive cells; Light grey, dotted line population= Myc negative cells; White, dashed grey line population = baseline. Representative of 3+ independent experiments with technical replicates. B) Quantification of the average median C3 fluorescence intensity from myc positive and myc negative cells within each sample. N=3 (1 experiment with 3 replicates; Representative of 3+ independent experiments) Statistics = t-tests. E) Average median C3 fluorescence intensities after normalization to the myc negative cells from each experimental group. ANOVA (p<0.001; F(4, 10)=74.47. N=3 (3 independent experiments). For all graphs * = p <0.05; ** = p<0.01 # = p<0.001 for all Myc + groups compared to M-CR2.

## Discussion

We have shown that truncated Sez6L2 inhibits cell lysis by the classical and alternative complement pathways. It does this by destroying active C3 convertases. Sez6L2 functions as a cofactor for Factor I to facilitate cleavage of C3b (but not C4b). This allows Sez6L2 to specifically disable the alternative pathway C3 convertase, C3bBb. Cleavage of C3b not only limits the initiation and amplification of the alternative pathway, but it likely also impairs C5 convertases that depend on C3b for their activity and ability to initiate the lytic pathway. Additionally, Sez6L2 deactivates C3 convertases by accelerating their decay which is defined as the dissociation of the active protease component from the complex. Sez6L2 removes C2a from the classical/lectin C3 convertase C4b2a and Bb from the alternative pathway C3 convertase C3bBb. Furthermore, we have shown that the complement inhibitory activity is shared within the Sez6 family. Full-length Sez6, Sez6L, and Sez6L2 all inhibit C3 deposition by the classical and alternative pathways. While most of these data do not employ neuronal specific paradigms, we have used well established methods of complement activity in serum to ascertain novel Sez6L2 protein functions that are still relevant to complement in the brain or other tissues and the molecular function of the Sez6 family.

Other labs have shown that Sez6 family proteins are known to control dendritic branching and synaptic density by an unknown mechanism [14, 16, 17, 25]. Additionally, genetic loss of the entire Sez6 family results in impaired motor coordination and motor learning, and impaired cognition [14–17]. Sez6L2 proteins can be found throughout the somatodendritic compartment in some neuronal cell types like Purkinje cells of the cerebellum [15, 16]. However, we report here that Sez6L2 can also be localized to synapses as we found in the CA1 region of the hippocampus. This synaptic location is also in line with a proteomic screen on dissociated cortical neurons reported by Loh et al. that found Sez6L2 within the excitatory synaptic cleft [60]. The synaptic and dendritic localization of Sez6L2 puts it in an ideal location to protect synapses and dendrites from complement-dependent pruning during development and may be a mechanism by which Sez6 proteins modulate synapse numbers and dendritic morphology. However, Sez6 proteins could also modulate complement activation levels affecting neurogenesis, neuronal migration, or immune reactions to various insults. Perhaps complement dysregulation explains the genetic association of the Sez6 family with multiple neurodevelopmental and psychiatric disorders including: autism, schizophrenia, intellectual disability, epilepsy, and bipolar disorder [1–9].

We found that both full-length and truncated versions of Sez6L2 were capable of inhibiting complement. This is likely important as the extracellular domains of all Sez6 family proteins can be cut near the transmembrane region by BACE enzymes (1 and 2) [17, 61,62]. This provides a means to remove the complement inhibitors from the cell surface making the cell vulnerable to complement deposition and its downstream consequences. On the other hand, it also provides a mechanism to release soluble complement inhibitors that could be useful to nearby “self” surfaces or the CSF. Soluble Sez6 is elevated in the CSF of patients with schizophrenia, bipolar, major depression and inflammatory pain compared to controls [63, 64]. Yet Sez6 in the CSF is decreased in Alzheimer’s [65]. Thus, altered shedding of the extracellular complement regulatory region of Sez6 proteins is likely connected and may have direct consequences on development of neuropathology. Cells may also change their vulnerability to complement by expressing alternative short splice variants of Sez6 family members that do not contain a transmembrane anchor and the three tandem CCP domains that presumably mediate complement inhibition [14, 16].

The domain structure of the Sez6 family which contains five CCP domains (three of which are consecutive) initially prompted us to investigate the complement regulatory functions of Sez6 family members. Although proteins with multiple CCP domains are common to the complement pathway, there are many other proteins that contain CCP domains (even tandem CCP domains) that do not have complement inhibitory functions. Although the amino acid sequences of CCP domains vary considerably, RCA proteins bind to C3b using a common binding platform on C3b with their 3-4 consecutive CCP domains in an extended orientation [32]. Ojha et al recently used AI-assisted computer learning along with significant functional annotation based off of decades of experimental research on RCA proteins to propose five short motifs that potentially confer complement inhibitory activity when conserved in a specific order across three tandem CCP domains [30]. Using their web-based CoreDo program, we found Sez6L2 has the correct motif pattern for predicted complement regulatory activity (Supplemental Figure 3; [30]). Thus, our results on Sez6L2 match their prediction. However, Sez6 and Sez6L do not have the correct five motif pattern; and yet, Sez6 was even more efficient at inhibiting C3b/C3c deposition by the classical pathway than Sez6L2. Interestingly, CSMD1, another brain expressed complement inhibitor associated with neurodevelopmental disorders, and perhaps a distant cousin of the Sez6 family, was also one of only two experimentally validated complement inhibitors that did not fit the five motif pattern identified and reported by Ojha et al [30, 66–69]. As the core CCP motif pattern is somewhat different between Sez6L2 and Sez6, it is possible that there are unique features to their complement regulatory activity yet to be discovered. In this study, we focused on the CCP domains and their C3 convertase inhibitory activity. Yet, the CUB domains of Sez6 proteins may also mediate more interactions with complement factors in order to boost, broaden, or limit their impact on the complement pathways. Interestingly, different splice variants of the Sez6 family also yield proteins with two or three CUB domains. The Sez6L variant we tested contained three CUB domains and was the least effective family member at blocking C3 deposition by the classical pathway. In contrast, the variants of Sez6 and Sez6L2 we tested had only two CUB domains and were much more effective at blocking the classical pathway. Further studies will investigate whether the CUB domains modulate the complement inhibitory activity of the Sez6 family members or whether differences in their activity are intrinsic to CCP domains.

The complement inhibitory activity of the Sez6 family may also explain their role in cancer. Increased expression of Sez6 family members has been linked to increased tumor growth and a poor prognosis in various cancers [18–24]. The role of complement in tumorigenesis is complex as it may exacerbate or inhibit tumor growth depending on the immune response and inflammatory environment. However, increased expression of complement regulatory proteins has been repeatedly shown to limit innate immune surveillance and cytotoxicity by the complement system and provide resistance to antibody based immunotherapies [70]. Increased expression of complement regulators may also dampen the B and T cell immune response to tumor cells after chemotherapy due to complement/immune cell crosstalk mediated via CR2 [70, 71]. Thus complement resistance not only influences the course of the disease but also limits therapeutic options. Testing for increased Sez6 family expression and employing strategies to block their complement inhibitory function alongside other therapeutic approaches may be necessary as it has been against tumors overexpressing other complement regulators like MCP, DAF, Factor H, or CD59.

## Materials and Methods

### Purified proteins

#### Cloning and Plasmids

The expression plasmid for Sez6L2-MH, a truncated version of human Sez6L2 that has the transmembrane and cytoplasmic region replaced by a tandem Myc/6x-His tag, was cloned using PCR from the Sez6L2 NCBI Refseq BC000567 cDNA (similar to <u>NM 001114100.2;</u> UniprotACC: Q6UXD5-6;) and the following primers: 5’-GACTCGAGAATTCGCGGCCGCCACCATGGGGACTCCCAGGGCCCAGCA-3’ and 5’-TTCTAGAAAGCTTGGTACCTCCCCCCCTTCCAGCTGCCGTGATGG-3’. The PCR product was digested with restriction enzymes Not1 and HindIII and cloned into the mammalian expression plasmid, pEZYmyc-His (Addgene plasmid #18701; http://n2t.net/addgene:18702; RRID: Addgene_18701; a gift from Yu-Zhu Zhang [72]), that was digested with the same enzymes. The human His-DAF plasmid from Sinobiological (HG10101-NH; modified from NCBI RefSeq NM_000574.3) was also used for protein production of H-DAF.

#### His-Tagged Protein Purification (Sez6L2-MH and H-DAF)

FreeStyle 293-F cells grown in serum-free FreeStyle 293 Expression Medium (Gibco, 12338-018) were transfected with Sez6L2MH or His-DAF plasmids using Fectopro (Polyplus, 116-001). After 48-72 hours, media was collected and His tagged proteins were bound using HisPur Ni-NTA Resin (ThermoFisher; 88221) in the presence of Calbiochem’s EDTA-free protease inhibitor cocktail 1:1000 (MilliporeSigma; Set V; 539137). Bound proteins were eluted in 250mM imidazole in PBS pH 7.4 and filtered through a 0.22um membrane (such as MilliporeSigma, SCGP00525). Elutions were then either dialyzed with PBS overnight (MWCO 12-14K, Spectrum Spectra/Por Molecular porous tubing, 132706) and concentrated using Amicon Ultra filters MWCO-10K or 30K (MilliporeSigma; 10K: UFC901024, 30K: UFC903024) or directly concentrated and put through multiple rounds of buffer exchange to PBS using the Amicon Ultra filters or similar product (Pierce 30K MWCO concentrator, 88529). If protein concentration resulted in a few visible protein aggregates, the solution was again filtered through a 0.22um membrane (such as MilliporeSigma Millex-GV4 SLGVL0405). Protein concentration was determined by absorbance at 280nm with 1 A280 unit=1mg/ml for H-DAF and a 1% extinction coefficient correction (E1%) of 11.4 for Sez6L2-MH (calculated based off amino acid sequence using ProtParam). Aliquots were frozen and stored at −80°C.

#### Other Purified proteins

The negative control protein, bovine serum albumin or BSA, was purchased from MilliporeSigma (A3294). The lyophilized BSA was dissolved in PBS with 250mM imidazole then concentrated, exchanged to plain PBS, and stored similar to Sez6L2-MH and H-DAF. All other purified proteins were purchased from CompTech: C1s-enzyme(A104), C2(A112), C3 (A113); C3b (A114), iC3b (A115), C3d (A117), C4 (A105), C4b (A108), C4bBP (A109), Factor B (A135), Factor D (A136), Factor H (A137), Factor I (A138). These proteins were purified from human serum and are in PBS.

### Hemolytic Assay

#### Classical pathway

Complement mediated lysis of erythrocytes releases hemoglobin that can be measured by spectrophotometers at 415 nm. Antibody-sensitized sheep erythrocytes (EA; CompTech; B200) were rinsed and diluted in GVB^++^ buffer (0.1% gelatin, 5 mM Veronal, 145 mM NaCl, 0.15 mM CaCl_2_ 0.5 mM MgCl_2_. 0.025 % NaN_3_, pH 7.3) 400ug/ml of Sez6L2-MH or other purified proteins (in PBS) were pre-incubated at in a v-bottom 96 well plate for 15 minutes on ice with 0.4% normal human serum (NHS, CompTech) in a final 90μl buffer solution equivalent to 40% GVB^++^ and 50% PBS with 0.15 mM CaCl_2_ and 0.5 mM MgCl_2_ (PBS^++^). Next, 10μl of EAs (4×10^8^ cells/mL in GVB^++^) were added to each sample and the plate was incubated at 37°C for 30 minutes. During this incubation, cells were suspended with a multichannel pipet every 10 minutes. The reaction was stopped by adding 100ul of cold GVBE buffer (0.1% gelatin, 5 mM Veronal, 145 mM NaCl, 0.025 % NaN_3_, 10 mM EDTA, pH 7.3). Remaining erythrocytes were pelleted by centrifugation at 500xg for 3 minutes. 150ul of the supernatant was transferred to a flat-bottom 96-well plate and measured for absorbance at 415nm using a microtiter plate reader (SpectraMax M5). The absorbance obtained with just buffer, NHS, and EAs (no added complement inhibitory proteins) was set at 100% lysis and the absorbance obtained without serum added or when GVBE was used in place of GVB^++^ to inhibit complement was set at 0% lysis. Absorbance from full lysis in each experiment was also determined by replacing PBS with H_2_0 (at 50% final sample volume). Prior to testing the efficacy of complement inhibitors, the amount of each lot of normal human serum necessary to lyse each batch of EAs to 80-90% of the absorbance obtained by H_2_0 samples was determined by testing a range of NHS concentrations from 0-2%. These titrations were done to ensure that inhibitors were not overwhelmed by saturating levels of complement activity. The titrations revealed that 80-90% lysis was usually obtained with 0.4% serum (Supplemental Figure 1A). Supernatants were also analyzed by western blotting for C3 cleavage products.

#### Alternative Pathway

The alternative pathway hemolytic assays were performed similar to the classical pathway assay except that plain rabbit erythrocytes (Er, CompTech) were rinsed and diluted in GVB° (0.1% gelatin, 5 mM Veronal, 145 mM NaCl, 0.025 % NaN_3_, pH 7.3). 400ug/ml Sez6L2-MH or other purified proteins (in PBS) were pre-incubated for 15 minutes on ice with 7-10% normal human serum in a final 95μl buffer solution containing 12.5mM MgCl_2_ and 12.5mM EGTA and the equivalence of 45% GVB° and 50% PBS. Next, 5μl of Ers (5×10^8^ cells/mL in GVB°) were added to each sample and the plate was incubated at 37°C for 30 minutes and processed similar to the classical hemolysis experiments. For alternative hemolysis assays, the NHS was titrated to lyse rabbit erythrocytes to ~30% of the absorbance (A415) obtained by full lysis by H_2_0 (Supplemental Figure 1B). The absorbance obtained with just buffer, NHS, and Ers (no purified complement inhibitor proteins added) was set at 100% lysis. As much higher serum concentrations are required to assay the alternative pathway than the classical pathway, setting this lower level of hemolysis at 100% allowed us to keep the serum concentration lower in order to detect inhibition by purified complement regulators.

### Cleavage of C3b or C4b by Factor I

C3b (4μg) or C4b (4μg) was incubated with 0.5μg Factor I alone or in the presence of 1 μg FH, 1μg C4BP, or 1-10μg Sez6L2 in PBS^++^ (Total volume ~30μl) for 2-8 hours at 37°C. The samples where then run on reducing SDS-PAGE gel and analyzed by western blot or imperial blue (coomassie) staining to identify C3 or C4 cleavage products.

### Decay Accelerating ELISA

#### Alternative C3 Convertase Decay Assay

An ELISA-based assay was used to measure decay accelerating activity towards the alternative pathway C3 convertase, C3bBb. ELISA plates (Nunc, MaxiSorp, 96 well plates, 44-2404-21) were coated with 100ul human C3b at 3 μg/mL in PBS (covered, 20 hours at room temperature). Wells were washed 2 times with PBS and blocked with 1% BSA in PBS for 30 minutes at room temperature. Wells were again washed twice with PBS. C3 convertase was formed by incubating with Factor B (0.5 μg/mL) and Factor D (50 ng/mL) in GVB° buffer with 3.5 mM NiCl_2_ at 37°C for 30 minutes. After, wells were washed 3 times with PBSt (0.1% tween). Factor H and Sez6L2-MH were mixed at various concentrations ranging from 0 μg/mL to 500 μg/mL in solutions with a final composition of 90 μL PBS to 225 μL GVB° with 3.5 mM NiCl_2_. 100ul of the Factor H or Sez6L2-MH mixtures were added to the plate in triplicate and incubated at 37°C for 45 minutes. Supernatants were collected for western blot analysis and the wells were washed 3 times with PBSt. Residual Bb on the plate was detected using polyclonal goat anti-Factor B (Comptech, A235, 1:4,000) for 1 hour at room temperature. Wells were washed 3 times with PBSt. Wells were then incubated with a donkey HRP-conjugated anti-Goat IgG (Azure Biosystems, AC2149, 1:10,000) for 45 minutes at room temperature. Wells were washed 3 times with PBSt. Color was developed using TMB substrate (ThermoScientific, N301) before adding 0.16 M sulfuric acid to stop the reaction. Absorbance was measured at 450 nm, and percent convertase remaining was calculated with the following formula = ((Ab450 inhibitor – Ab450 background)/(A450 buffer – Ab450 background)) x 100. Background values were obtained from wells not coated with C3b but incubated with Factor B and D. Nonlinear regression utilized a variable slope and four parameters.

#### Classical/Lectin C3 Convertase Decay Assay

This ELISA-based assay was performed similar to the C3bBb assay outlined above but with the following changes: Plates were coated with 3ug/mL C4b in PBS. After washing and blocking with 1% BSA, C3 convertases were formed by incubating with C2 (0.5-1ug/ml) and C1s-enzyme (0.3ng/mL) in GVB++ at 37 for 30 minutes. After, wells were washed 3 times with PBS^++^. H-DAF and Sez6L2-MH were mixed at various concentrations ranging from 0 μg/mL to 500 μg/mL in solutions with a final composition of 50% GVB^++^ and 50% PBS^++^. 100ul of the H-DAF or Sez6L2-MH mixtures were added to the plate in triplicate and incubated at 37°C for 30 minutes. Wells were washed 3 times with PBSt^++^ (containing 0.1% tween). Residual C2a/C2 on the plate was detected using polyclonal goat anti-C2 (Comptech, A212, 1:6,000 in PBSt^++^) for 45 minutes at room temperature. Wells were washed 3 times with PBSt^++^ and then incubated with a donkey HRP-conjugated anti-Goat IgG (Azure Biosystems, AC2149 1:10,000 in PBSt^++^) for 45 minutes at room temperature and then processed as above. Background values were obtained from wells not coated with C4b but incubated with C2 and C1s-enzyme.

### Flow Cytometry

CHO (Freestyle) cells were grown in serum-free Freestyle media (Gibco, 12651-014) and transfected with human cDNA expression plasmids with N-terminal Myc tags obtained from Sinobiological: M-SEZ6L2 (HG13969-NM; modified from NCBI RefSeq BC000567; similar to <u>NM 001114100.2;</u> Uniprot ACC: Q6UXD5-6;), M-SEZ6 (HG13436-NM; modified from NCBI RefSeq <u>NM 178860.4;</u> Uniprot ACC: Q53EL9-1;), M-SEZ6L (HG20982-NM; modified from <u>NM 001184773.1</u> and contains the point mutation 1428A/C that does not alter the amino acid sequence; Uniprot ACC: Q9BYH1-6), M-MCP (HG12239-NM; modified from NCBI RefSeq <u>BC030594</u>), or M-CR2 (HG10811-NM; modified from NCBI RefSeq <u>NM 001877.3</u>) using FectoPro transfection reagent (Polyplus, 116-001). Alternatively cells were transfected with a GFP expression plasmid or co-transfected with GFP and M-SEZ6L2 or HIS-tagged DAF (H-DAF; Sinobiological HG10101-NH; modified from NCBI RefSeq <u>NM 000574.3</u>). After 18-24 hours, the cells were collected and loaded at 4×10^5^ cells per well in a 96-well v-bottom plate. To assay the activity of the classical complement pathway, CHO cells were sensitized with a 30 minute incubation with rabbit anti-hamster lymphocyte serum (1:10; Cedarlane, CLA14940) in 100μl cold FACs buffer (0.5% BSA in PBS) at 37°C. Cells were washed twice with FACs buffer then incubated with 15% C5-depleted human serum (Comptech, A320) or a range of C5 depleted serum from 0-20% in GVB++ at 37°C for 60 minutes. After washing 2x with cold FACs buffer, cells were incubated with a mouse anti-human C3 APC conjugated antibody (1:50; Biolegend, 846106, clone 3E7iC3b) and mouse anti-Myc Alexa488 conjugated antibody (1:50; Cell Signaling Technologies, 2279S, clone 9B11) for 30 minutes at 4°C. For samples transfected with GFP, only the anti-C3 APC antibody was used. After 2 washes in FACs buffer, cells were resupended in FACS buffer with 1μl propidium iodide (PI, Invitrogen, BMS500PI) and processed using a BD Accuri C6 Flow Cytometer. Data was analyzed by FlowJo™ Software (Windows Version 10.6.1: Ashland, OR: Becton, Dickinson and Company; 2019). Data were gated for the single cell population (FSC-A:FSC-H) and PI negative population. To assay the alternative pathway, cells were processed as above except cells were sensitized with only a low level of anti-hamster lymphocyte serum (1:50) and the 20% C5 depleted serum was diluted in GVB° with 10mM EGTA and 10mM MgCl_2_ to block the classical pathway.

### Western blot

Supernatants from hemolysis assays, decay accelerating ELISA’s, or Factor I cleavage assays were mixed with SDS-loading dye and run on 8-15% SDS-PAGE gels and transferred to PVDF. Membranes were blocked with 5% milk in TBSt (tris buffered saline (20 mM Tris-Cl, pH 7.4; 150 mM NaCl) with 0.1% Tween 20) for 30 minutes and probed with primary antibodies: polyclonal goat anti-C3d (R&D systems, AF2655, 1:1000), polyclonal sheep anti-Sez6L2 (R&D Systems, AF4916; 1:2000), monoclonal mouse anti-myc ((Developmental Studies Hybridoma Bank; 9E10, 1:2000), polyclonal goat anti-Factor B (CompTech, A235, 1:1000), polyclonal goat anti-C4 (CompTech, A205; 1:1000) in 5% milk in TBSt for 1 hour at room temperature or overnight at 4°C. Membranes were washed 3 times in TBSt and then incubated with HRP secondary antibodies (anti-Mouse HRP, Biorad, 170-6516, 1:5,000; anti-Goat HRP Azure Biosystems, AC2149, 1:10,000; anti Sheep HRP, Jackson Labs, 713-035-003, 1:1000) for 1 hour at room temperature in 5% milk in TBSt. After washing we applied ECL substrate (Pierce) and developed membranes using a digital imager (Azure Biosystems). Membranes were occasionally stripped using buffer consisting of 200mM glycine; 0.1% SDS, 1% Tween 20, pH 2.2 and re-probed. Western blot band densities were quantitated using Image J/FIJI.

### Immunohistochemistry (IHC) and Imaging

Animal care and use were carried out in compliance with the US National Research Council’s Guide for the Care and Use of Laboratory Animals and the US Public Health Service’s Policy on Humane Care and Use of Laboratory Animals. Protocols were approved by the University Committee on Animal Resources at the University of Rochester. Sez6 Triple Knockout mice (Sez6 TKO) have a complete knockout of Sez6, Sez6L and Sez6L2 [15, 16]. They are also known as BSRP TKO mice and were obtained from Dr. Jenny Gunnersen, University of Melbourne; Australia. Mice were anesthetized with ketamine/xylazine (100 and 10 mg/kg, respectively) and intracardially perfused for 1 minute with phosphate-buffered saline (PBS) containing EDTA (1.5 mg/ml) followed by 4% paraformaldehyde (PFA) in PBS. Brains were post-fixed for 18-24 hours in 4% PFA then stored in PBS at 4°C. Brains were cut into 40μm-thick coronal using a vibratome (Leica V1000) and stored in a cryoprotectant mixture of 30% PEG300, 30% glycerol, 20% 0.1 M phosphate buffer, and 20% ddH_2_O at −20°C. IHC was performed on free-floating-sections. The sections were washed three times for 30 minutes in PBS to remove the cryoprotectant. Then sections were incubated in 100mM glycine in PBS for 30 minutes followed by citrate antigen unmasking at 37°C for 30 minutes (Vector, H3300 with 0.05% tween 20). Sections were blocked overnight in blocking buffer (consisting of 1.5% BSA (MilliporeSigma; A3294), 3% normal donkey serum (MilliporeSigma, 566460), 0.5% Triton X100 (Promega, H5142), and 1.8% NaCl in 1× PBS) with donkey anti-mouse Fab fragments (Jackson ImmunoResearch Laboratories, Cat# 715-007-003; 1:300). Sections were then washed three times in 1× PBS with 1.8% NaCl. Primary antibodies: polyclonal sheep anti-Sez6L2 (R&D Systems, AF4916,1:500) and polyclonal chicken anti-Homer1 (Synaptic Systems, 160006, 1:500)) were diluted in blocking buffer and incubated with brain sections for 1-3 days at room temperature with agitation. Sections were then washed three times for 30 min in 1× PBS with 1.8% NaCl and then incubated overnight at room temperature in Alexa Fluor-conjugated secondary antibodies (1:500 Jackson ImmunoResearch Laboratories Alexa 488 Donkey anti-sheep, Cat# 713-545-147, RRID: AB_2340745; and Alexa 647 Donkey anti-chicken, Cat# 703-605-155, RRID: AB_2340379) in blocking buffer. Finally, sections were washed 3 times with 1×PBS with 1.8% NaCl, mounted on slides with Prolong Diamond mounting agents with DAPI (Life Technologies; P36962).

IHC sections were imaged with a Hamamatsu ORCA-ER camera on an Olympus BX-51 upright microscope with Quioptic Optigrid optical sectioning hardware with a 10x, NA 0.4 or 60x, NA 1.4 objective with *z*-step = 0.3μm. Volocity 3DM software (Quorum Technologies; version 6.3) was used for image collection and analysis. Differences in *z*-axis registration of various fluors were corrected by calibration with multicolor fluorescent beads. Image stacks are displayed as maximum intensity protections.

### Statistics

GraphPad Prism software version 8.3.0 for Windows (La Jolla California USA) was used to perform all statistics. Statistical analysis was generally performed with 1-way ANOVAs followed by Holm-Sidak’s multiple comparisons test (MCT). For the CHO cell, C3 deposition assays Myc positive and Myc negative populations within the same sample were compared with multiple t tests and p values were corrected with Holm’s-Sidak’s MCT for the many samples within the experiment. N values for each experiment are specifically stated and defined in the figure legends. We defined significance as p <0.05 and used the following markings on graphs * p<0.05; ** p<0.01. All data are expressed as the mean ± standard error of the mean (SEM).

## Author Contributions

JWH designed the study. WQQ, SL, and JWH did the hemolysis assays. SL, SAM, and JWH did the factor I cleavage assays, SAM and JWH did the decay accelerating assays. WQQ, PS, and JWH did the CHO cell/C3 deposition flow cytometry assay. JWH did the C3-SEZ6L2 interaction ELISA. JWH and HL performed the IHC. JWH wrote the manuscript. JG provided expertise on the SEZ6 family. All authors discussed, edited, and approved the manuscript.

## Acknowledgements

This work was supported by funding from the National Institutes of Health (R21NS111255), the Harry T. Mangurian Jr. Foundation, the National Health and Medical Research Council (NHMRC, GNT1099930 and GNT1140050), and the Judith Jane Mason and Harold Williams Memorial Foundation.

## Supplemental Figures

**Supplemental Figure 1:**
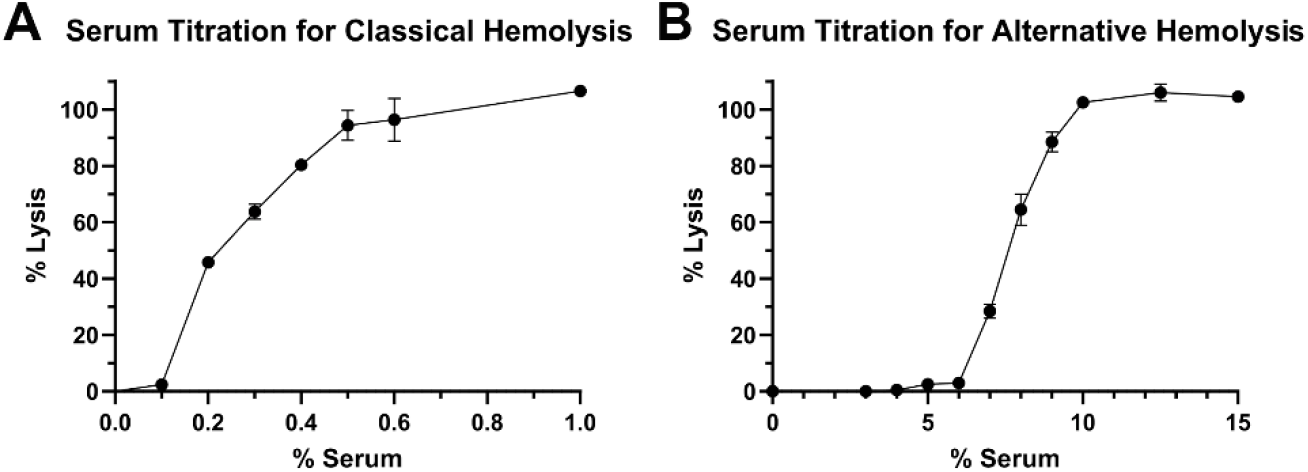
Serum titrations for hemolysis assays. Human serum was added at increasing concentrations in classical or alternative hemolysis assays to determine the amount necessary to yield lysis of erythrocytes.

**Supplemental Figure 2:**
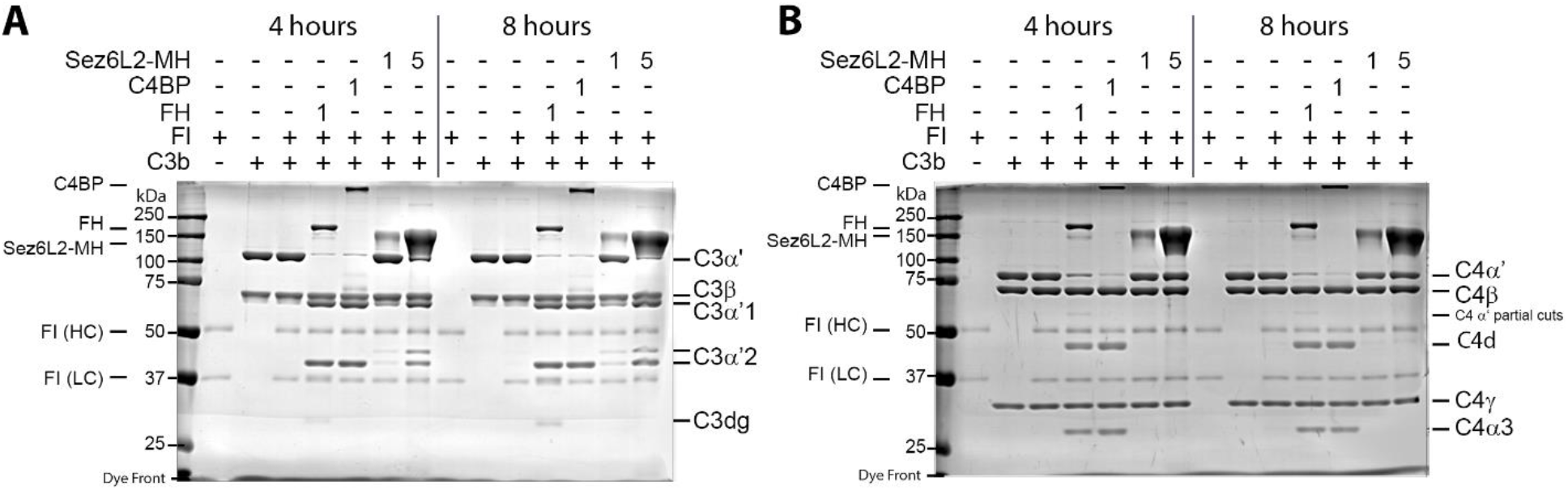
Factor I cleavage of C3b or C4b with Sez6L2-MH (4 and 8 hour experiments). Factor I cleavage assay of C3b (A) or C4b (B). C3b or C4b and Factor I (FI) were incubated alone, with Factor H (FH, 1ug), C4BP (1ug), or Sez6L2-MH (1ug or 5ug) for 4 or 8 hours at 37°C. Then samples were analyzed by coomassie stained gels. Sez6L2-MH aids significant factor I cleavage of C3b but not C4b at 4 and 8 hours resulting in appearance of the C3b cleavage products, C3α’1 and C3α’2, but not the C4b cleavage products, C4d and C4α3.

**Supplemental Figure 3:**
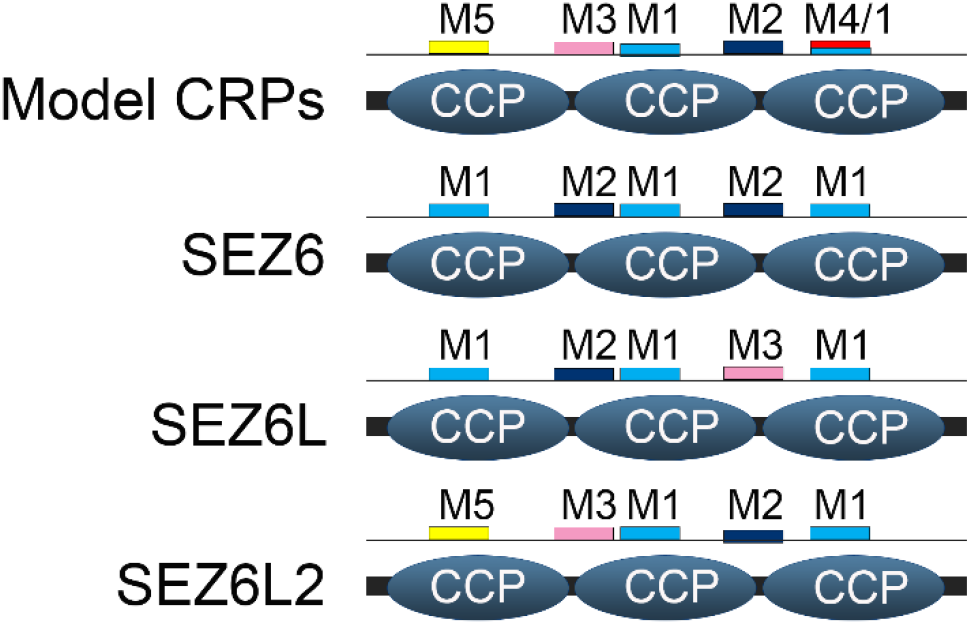
Sez6L2, but not Sez6 or Sez6L, contains the CCP motifs described by Ojha et al [30] in the correct order to predict complement regulatory function. Sez6 family FASTAs sequences were uploaded into the web-based program provided by Ojha et al (http://coredo.nccs.res.in/meme-5.0.3/CoReDo/home.html). Classical model complement regulatory proteins (CRPs) have motifs in the order of M5, M3, M1, M2, and either M4 or M1 spaced across 3 consecutive CCP domains. Sez6L2 has this same pattern, but Sez6 and Sez6L do not.

